# A Mathematical Model of the Phosphoinositide Pathway

**DOI:** 10.1101/182634

**Authors:** Daniel V Olivença, Inna Uliyakina, Luis L Fonseca, Margarida D Amaral, Eberhard Voit, Francisco R Pinto

**Author notes:** Corresponding Author Contacts (ext. 28232).

## Abstract

Phosphoinositides are signaling lipids that constitute a complex network regulating many cellular processes. We propose a computational model that accounts for all known species of phosphoinositides in the plasma membrane of mammalian cells. The model replicates the steady-state of the phosphoinositide pathway and most known dynamic phenomena. Furthermore, sensitivity analysis demonstrates model robustness to moderate alterations in any of the parameters. Model analysis suggest that the greatest contributor to PI(4,5)P_2_ production is a flux representing the direct transformation of PI into PI(4,5)P_2_ and is also responsible for the maintenance of this pool when PI(4)P is decreased. PI(5)P is also shown to be a significant source for PI(4,5)P_2_ production. The model was validated with data from siRNA screens that knocked down the expression of several enzymes in the pathway. The screen monitored the activity of the epithelium sodium channel, ENaC, which is activated by PI(4,5)P_2_. Moderating ENaC activity can have a therapeutic effect in Cystic Fibrosis (CF) patients. Our model suggests control strategies where the activities of the enzyme PIP5KI or the PI4K+PIP5KI+DVL protein complex are decreased and cause an efficacious reduction in PI(4,5)P_2_ levels while avoiding undesirable alterations in other phosphoinositide pools.

**Abbreviations:**

AKT: Protein Kinase B, a serine/threonine-specific protein kinase
ASL: Airway surface liquid
BST: Biochemical systems theory
CF: Cystic fibrosis
DAG: Diacylglycerol
ENaC: Epithelial Sodium Channel
ER: Endoplasmic Reticulum
GMA: Generalized mass action
INPP5: Inositol polyphosphate 5-phosphatases
IP_3_: Inositol triphosphate
LTPs: lipid transport proteins
MCSs: membrane contact sites
MDCK cells: Madin-Darby Canine Kidney Epithelial Cells
MTM: Myotubularin
OCRL: Lowe Oculocerebrorenal Syndrome Protein; OCRL is an INPP5
ODE: Ordinary differential equations
PI: Phosphatidylinositol
PI(3)P: phosphatidylinositol 3-phosphate
PI(3,4)P_2_: Phosphatidylinositol 3,4-biphosphate
PI(3,4,5)P_3_: phosphatidylinositol 3,4,5-triphosphate, with phosphates in the third, fourth and fifth positions
PI(3,5)P_2_: Phosphatidylinositol 3,5-biphosphate
PI(4)P: phosphatidylinositol 4-phosphate
PI(4,5)P_2_: Phosphatidylinositol 4,5-biphosphate with phosphates in the fourth and fifth positions of the inositol ring
PI(5)P: phosphatidylinositol 5-phosphate
PI3K: Phosphoinositide 3-kinase
PI4K: Phosphoinositide 4-kinase
PIP5K: Phosphoinositide 4-phosphate 5-kinase
PIKfyve: FYVE finger-containing phosphoinositide kinase.
PLC: Phospholipase C
PLD: Phospholipase D
PLIP: PTEN-like lipid phosphatase
PTEN: Phosphatase and tensin homolog
PKC: Protein kinase C
SAC: Suppressor of actin
SHIP1: SH2 domain-containing phosphatidylinositol 5’-phosphatase
SKIP: Skeletal muscle and kidney enriched inositol polyphosphate phosphatase
SYNJ: Synaptojanins
TPIP: PTEN-Like Inositol Lipid Phosphatase
Wnt3a: Wingless-Type MMTV Integration Site Family, Member 3A
DVL: Segment Polarity Protein Dishevelled Homolog DVL

## 1. INTRODUCTION

Biological systems have evolved by improving the efficiency of complex regulatory networks that control multiple mechanisms in the cell through the fine balancing of the same enzymatic reactions. Phosphoinositides are important lipids interconverting into each other by multiple enzymatic reactions which altogether constitute an example of such a complex network which regulates most cellular functions. Phosphoinositides are key signaling messengers and several play important parts in regulating physiological processes including vesicular trafficking, transmembrane signaling, ion channel regulation, lipid homeostais, cytokinesis and organelle identity as characteristic identifiers for different membranes in the cell ^1–5^. It is thus not surprising that phosphoinositides play critical roles in a number of pathological conditions including immunological defense, mediating replication of a number of pathogenic RNA viruses, in the development of the parasite responsible for malaria, in tumorigenesis, Alzheimer’s disease, diabetes, etc ^6–9^.

Their inositol head can be phosphorylated at its third, fourth and fifth carbons thus creating different subspecies. The phosphoinositide pathway connects eight metabolites through a dense network of 21 chemical reactions, which are catalyzed by 19 kinases and 28 phosphatases ^10^ (Figure 1). The resulting degree of complexity prevents simple interpretations and renders intuitive predictions of pathway behavior and regulation unreliable. It is especially difficult to pinpoint the roles of less abundant phosphoinositides, such as PI(5)P and PI(3,4)P_2_. PI(4,5)P_2_ is present throughout the plasma membrane and considered a general marker for the cell membrane. By contrast, PI(3,4,5)P_3_, marks the basolateral part of a polarized cell’s membrane but is absent from the apical part ^1,11^.

**Figure 1.**
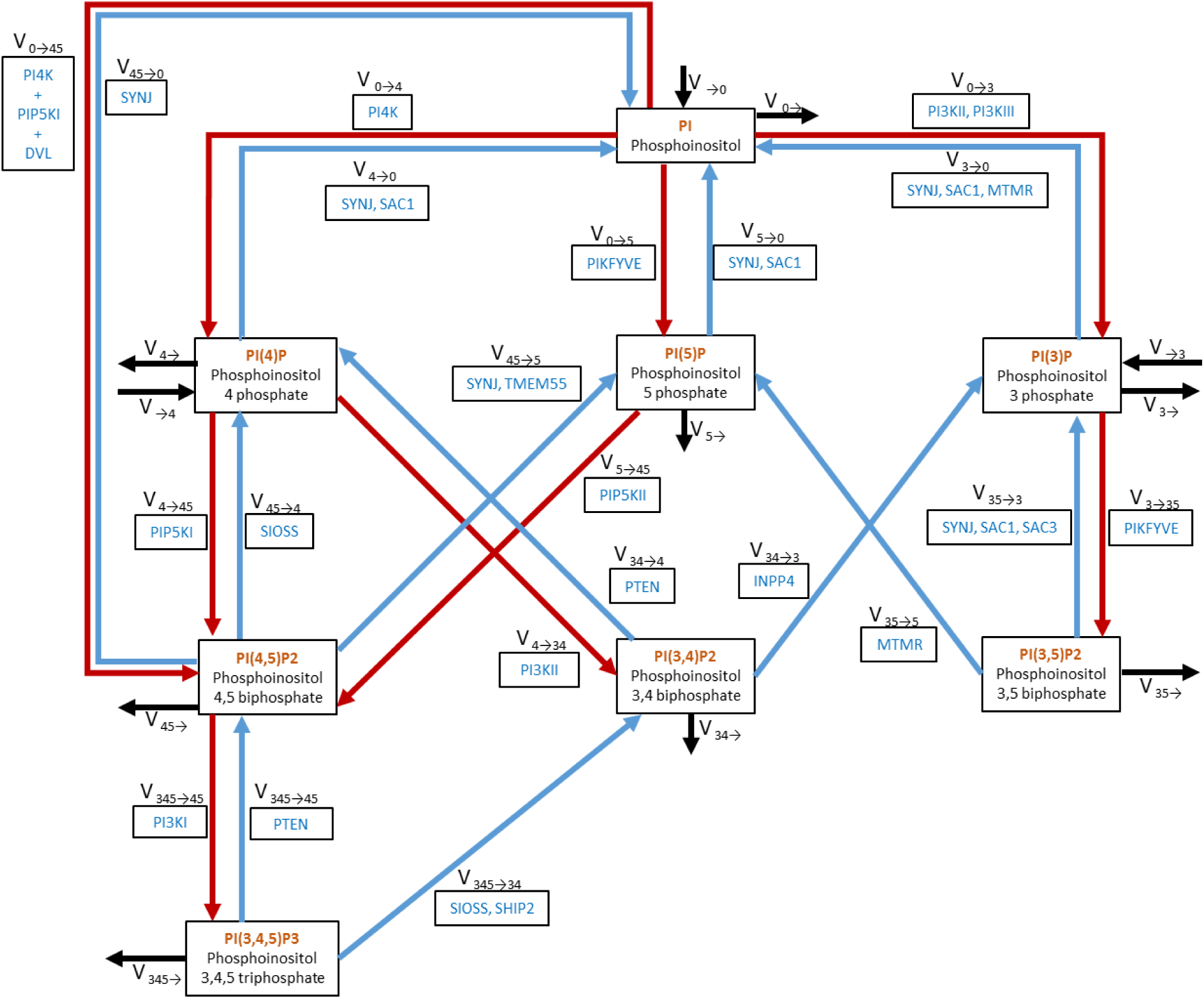
Map of the phosphoinositide pathway. Red arrows represent fluxes of phosphorylation and blue arrows fluxes of hydrolysis. For each flux, the name (v_i→j_) and the group of enzymes that catalyze the reaction are shown. Black arrows represent input and output fluxes of material entering and leaving the system. SIOSS is a group of phosphatases, consisting of SYNJ 1/2, INPP5 B/J/E, OCRL1, SAC2 and SKIP (skeletal muscle and kidney enriched inositol polyphosphate phosphatase). PI4K+PIP5KI+DVL denotes a complex formed by the three proteins. Proteins separated by commas catalyze the same reaction. INPP5: Inositol polyphosphate 5-phosphatases; OCRL1: Lowe Oculocerebrorenal Syndrome Protein.

Other phosphoinositides characterize intracellular membranes (Figure 2). PI(3,5)P_2_ is typical for multivesicular bodies and lysosomes, whereas PI(4)P is found in the Golgi and phosphatidylinositol (PI) is located in the ER ^12^. To achieve this distinctive variability in phosphoinositides composition among different membrane compartments, the cell must be able to modulate phosphoinositides metabolism in a targeted, localized manner.

**Figure 2.**
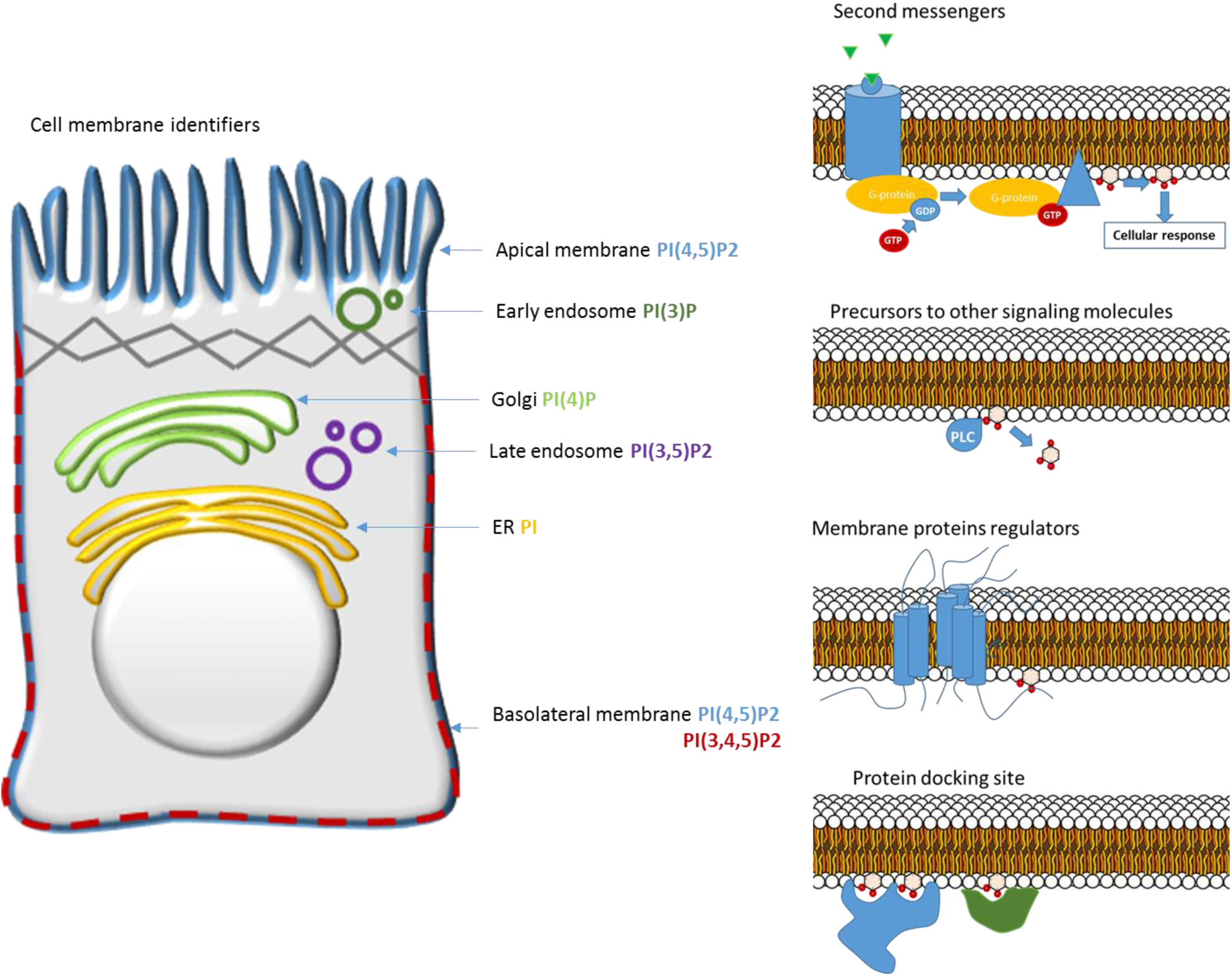
Functions of the phosphoinositides in the cell. Phosphoinositides are signaling lipids that are cell membrane identifiers. PI(4,5)P2 marks the plasma membrane, PI(3)P the early endosomes, PI(4)P the Golgi, PI(3,5)P2 the late endosomes, PI the ER and finally PI(3,4,5)P3 is present in the basolateral part of the plasma membrane and absent from the apical part. Phosphoinositides are also second messengers, precursors to other signaling molecules and membrane protein docking sites and regulators.

Phosphoinositides also serve as precursors for various signaling molecules, as in the case of PI(4,5)P_2_, which can be transformed into diacylglycerol (DAG) and inositol triphosphate (IP3) through the action of phospholipase C (PLC). They are furthermore docking sites in the plasma membrane, for instance, for AKT (also known as Protein Kinase B) in the case of PI(3,4,5)P_3_.

Interestingly, phosphoinositides are also key regulators of ion channel activity ^11^. The epithelial sodium channel (ENaC) is of interest, as it plays a critical role in Cystic Fibrosis (CF), a genetic condition caused by mutations in the gene encoding CFTR, a chloride channel that also regulates other ion conductance, namely through ENaC, across epithelia. In order to keep ENaC open, lysine residues present at the N terminus of the β and δ subunits need to be bound to PI(4,5)P_2_ ^13^. This channel function is upregulated in the lungs of individuals with CF, and the increased absorption of sodium and water is considered to be the major cause of lung disease due to critical dehydration of airway surface liquid (ASL) ^14^. The dehydrated ASL and consequent impairment of mucociliary clearance, in turn, is a major cause of respiratory problems in CF ^13^. Thus, a better understanding of the phosphoinositide pathway is of paramount importance, as it may contribute to ameliorating the CF phenotype by manipulating the levels of PI(4,5)P_2_, which moderate the action of ENaC.

To address the challenges of complexity, it is advantageous to resort to mathematical models, which indeed have already been proposed for particular components of the phosphoinositide pathway. Narang ^12^, Xu ^15^, Nishioka ^16^ and Purvis ^17^ proposed models mainly focused on understanding the dynamics of PI(4,5)P_2_, PLC, IP3 and DAG, since these molecules are directly associated with calcium release and protein kinase C (PKC) activation, which are important signaling events. Other models, such as those developed by Araia ^18^ and MacNamara ^19^, focus on PI(4,5)P_2_, PI(3,4,5)P_3_, phosphoinositide 3-kinase (PI3K), phosphatase and tensin homolog (PTEN) and their roles in cancer. None of these models accounts for all phosphoinositide species. However, the inclusion of less abundant species is important for understanding the distinctions between membrane compartments and for rationalizing the observed impact of several enzyme knock-downs on PI(4,5)P_2_-mediated ENaC modulation ^13^.

Here, we propose a mathematical model of the complete phosphoinositide pathway. Our primary goal is to shed light on the dynamics of this pathway. Moreover, the model will facilitate a deeper understanding of the unique composition of membranes in different compartments and thereby provide an effective tool for exploring possible therapeutic targets for CF, cancer and other diseases in which the phosphoinositide pathway plays a critical role.

## 2. RESULTS

The primary result of this study is a mathematical model of the phosphoinositide pathway that contains all its molecular components and captures features of the pathway documented in the literature. The pathway map underlying the model is exhibited in Figure 1. Its structure is based on a review by Balla ^11^, but expanded with information from other sources ^2,3,10^. To facilitate the presentation and discussion of results, each flux is represented by v_i→j_, and the group of enzymes catalyzing it by E_i→j_, where i and j identify the phosphorylated positions of the substrate and product phosphoinositide species, respectively.

### 2.1. Consistency of the Model with Data

As described in the *Methods* section, model equations were formulated according to Biochemical Systems Theory (BST) ^20,21^. Initial parameter estimates were derived from the literature and from the BRENDA database ^22^. The parameter values were subsequently optimized with a genetic algorithm such that the model matched reported phosphoinositide steady-state levels (Supplementary Table ST1) and dynamic phenomena reported in the literature (Figure 3 a,c and Supplementary Table ST2). This model successfully mimics steady-state levels and 11 out of 13 observed phenomena. The observations that were not replicated are: 1) when the PI levels are reduced, the drop in PI(4,5)P_2_ levels is not as evident as reported in the literature (Figure 3a) and 2) the knockout of myotubularin MTMR2 effects are only partially replicated (Figure 3c).

**Figure 3.**
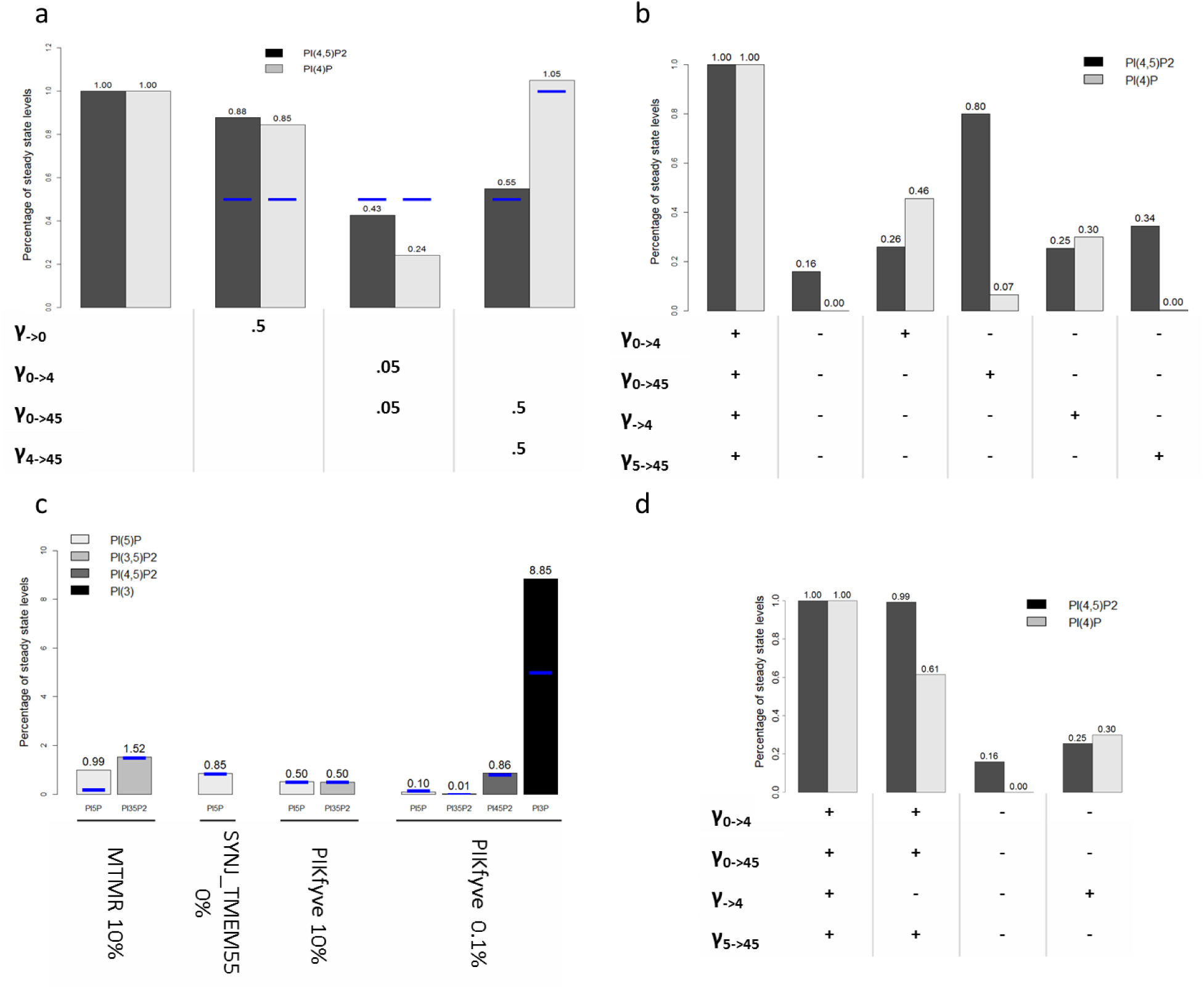
Perturbations to the phosphoinositide pathway. Blue lines represent experimental observations and bars represent model predictions. **a)** Perturbation of PI levels, PI4K and PI5KI activities and resulting effects in PI(4,5)P_2_ and PI(4)P. γ _→0_ is decreased to 50% to trigger a decrease of 50% in PI. **b)** Perturbation of input fluxes to the levels of PI(4)P and PI(4,5)P_2_. After stopping all inputs into PI(4)P and PI(4,5)P_2_, the inputs are re-activated, one at a time, to test if they are sufficient to restore PI(4,5)P_2_ levels. Enzyme knockouts were simulated by setting the rate constant of the corresponding flux to zero, except for γ_0→4_, which was decreased to 20% of its original value, in order to avoid numerical errors in the simulation due to very small levels of PI(4)P. **c)** Perturbations to MTMR, SYNJ_TMEM55 and PIKfyve that were used to fit the model to the behavior of phosphoinositides with small pools: PI5P, PI(3,5)P_2_ and PI(3)P. **d)** Consequences of Golgi PI(4)P input (γ _→4_) for the levels of PI(4)P and PI(4,5)P_2_ pools. Golgi PI(4)P has a significant impact on the PI(4)P pool but barely affects the PI(4,5)P_2_ pool.

### 2.2. Model sensitivities

The profile of model sensitivities is a double-edged sword. On the one hand, high sensitivities make the model susceptible to unreasonable responses from small perturbations or noise. On the other hand, if the system has a signaling function, small signals must be amplified to have appropriate effects. The model presented here has a stable steady-state that is mostly insensitive to parameter changes (Supplementary Table ST8). In fact, the system is even robust to large changes in parameter values (Supplementary Figure S1a, b and c). At the same time, the model does exhibits clusters of high sensitivities that are associated with signaling compounds (Figure 4).

**Figure 4.**
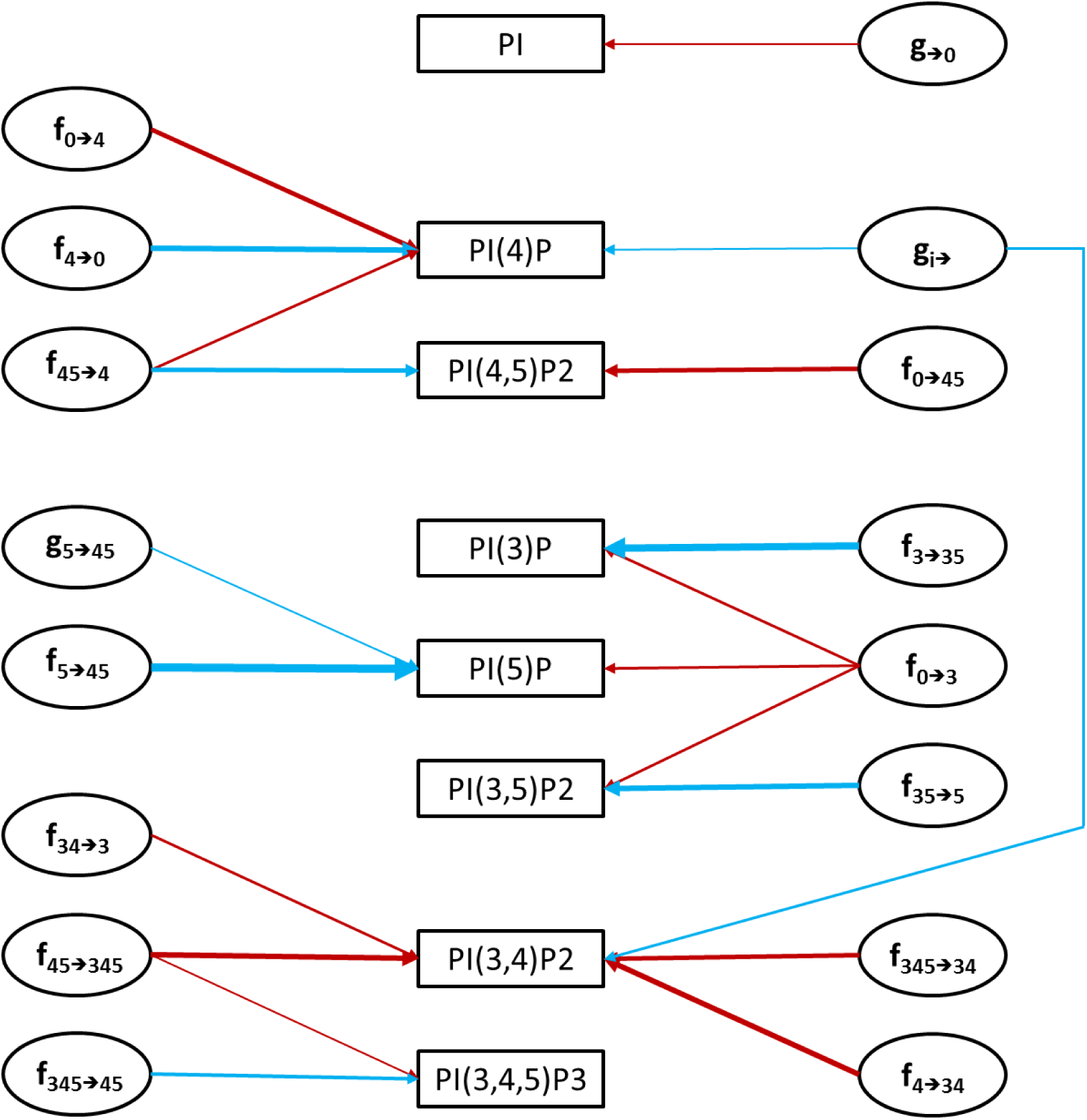
High-sensitivity network. Arrows represent amplifying sensitivities with absolute magnitude greater than 1. Red and blue arrows represent positive and negative sensitivities, respectively. The thickness of the arrows is proportional to the intensity of the sensitivities.

#### 2.2.1. Analysis of low sensitivities and parameter identifiability

Even though most sensitivities are low, one must question how many of the parameters are actually identifiable. To address this question, we performed a Monte Carlo search of the parameter space, which revealed that only 166 out of 79,993 parameter sets tested yield correct steady-state levels (less than 0.15%), given an acceptable material influx into the pathway. All of these 166 solutions have a worst adjustment score than the manually fitted set and the set found with a genetic algorithm (Figure 5). These results suggest that the model parameterization is sufficiently specific given the available experimental information.

**Figure 5.**
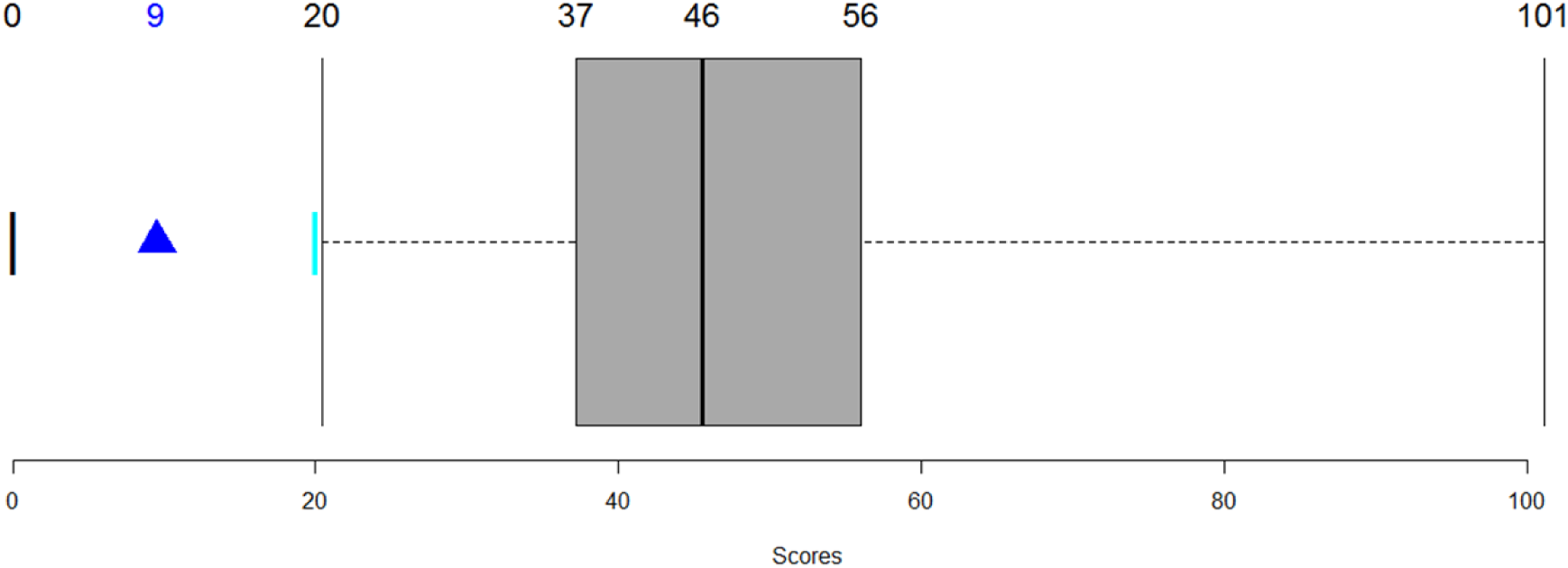
Sum of squared errors for parameter sets detected through Monte-Carlo parameter space exploration. All parameter sets shown comply with the following conditions: 1. phosphoinositide steady state levels are within the intervals retrieved from the literature; 2. the relative amounts between the phosphoinositide pools match the data; 3. influxes are less than 25% of the corresponding phosphoinositide pools; 4. effluxes are less than 7% of the corresponding phosphoinositide pools. The black bar at zero represent the score of a perfect model, the best set found by the genetic algorithm is shown as a blue triangle and the manually found set is the cyan bar. The boxplot concerns the 116 admissible alternative parameter sets.

#### 2.2.2. High-sensitivity sub-networks

The pairs of model variables and parameters with high sensitivities (Figure 4) form a network that clusters into four groups around: 1) PI, which is the source of the phosphoinositides; 2) PI(4)P and PI(4,5)P_2_, which are responsible for plasma membrane identification and PI(4,5)P_2_ maintenance; 3) the small lipids pools (PI(3)P, PI(5)P and PI(3,5)P_2_); and 4) PI(3,4,5)P_3_ and its derivate PI(3,4)P_2_.

This high-sensitivity network is reflected in a map of parameters that are best poised to serve as “master regulators” for controlling the variables in the different groups. For example, an increase in the levels of PI(3,4,5)P_3_ and PI(3,4)P_2_ is most easily accomplished by altering the kinetic order in the flux V_45→345_. An increase in V_345→45_ elicits a reduction of PI(3,4,5)P_3_, which highlights the importance of PI3KI and PTEN for this part of the pathway. If simultaneous increases in the levels of the three phospholipids PI(3)P, PI(5)P and PI(3,5)P_2_ are required, a researcher should boost V_0→3_. As an alternative, he could decrease each phospholipid independently manipulating the respective consumption fluxes.

### 2.3. New Insights into the Phosphoinositide System

The model can be used to shed light on the control of the phosphoinositide pathway. Particularly pertinent insights are described in the following subsections.

#### 2.3.1. PI(4,5)P_*2*_ is sensitive to PI, PI4K and PIP5KI

Model simulations replicating reported experimental results demonstrate that PI(4,5)P_2_ is sensitive to the level of PI and to the activities of phosphoinositide 4-kinase (PI4K) and phosphoinositide 4-phosphate 5-kinase (PIP5KI) (Figure 3a).

##### PI4K controls PI(4,5)P_*2*_ levels

According to the literature, a knockout of PI4K leads to a decrease in PI(4)P and PI(4,5)P_2_ to 50% of their basal level ^23^. Decreasing PI4K will cause not only the decrease of v_0→4_ but also V_0→45_ because this kinase is part of the protein complex that catalyzes V_0→45_. The model mimics this phenomenon for PI(4,5)P_2_ although it predicts a more severe drop in the levels of PI(4)P.

##### PIP5KI controls PI(4,5)P_*2*_ levels

One strategy for reducing PI(4,5)P_2_ levels is to decrease the amount of PIP5KI. This mechanism is probably viable *in vivo* because a single allele of the PIP5KIγ gene is sufficient to sustain life in mice embryos, whereas knock-out PIP5KIγ mice die shortly after birth ^24^. The same study also showed that *α* and *β* genes are not necessary to maintain viability, and their roles are still unclear. Volpicelli-Daley *et al.* ^24^ furthermore reported that PI(4,5)P_2_ levels drop around 50% in PIP5KIγ KO mice. Decreasing the activities of PIP5KI (E_4→45_) and PI4K/PIP5KI (E_0→45_) to 50% in the model, reduces PI(4,5)P_2_ to roughly 50% of its basal level.

##### PI controls PI(4,5)P_*2*_ levels

Kim ^25^ reported that a 50% drop in the PI pool causes a similar decrease in PI(4,5)P_2_ levels. PI(4,5)P_2_ in the model is sensitive to a reduction in PI but does not drop as much as reported in the literature. Specifically, a 50% drop in PI will only lead to a reduction of 11% in PI(4,5)P_2_. A 50% drop of PI in the whole cell would also affect other membrane compartments responsible for the production of PI(3)P and PI(4)P. To include this effect, we closed v→_4_ and v→_3_. However, this intervention decreases PI(4,5)P_2_ only to 88% of its basal level. Interestingly, PI(4)P drops to 47%. To achieve a 50% drop in the PI(4,5)P_2_ pool we would have to shut down v→_4_ and v→_3_ completely and reduce the influx of PI,v→_0_, to 2% of its original value.

#### 2.3.2. PI(3,4,5)P_*3*_ levels are sensitive to the concentrations of PTEN and PI3KI

PTEN has been known to be a tumor suppressor for almost twenty years ^11^. This phosphatase hydrolyzes the third position of the phosphoinositide inositol ring in PI(3,4,5)P_3_ into PI(4,5)P_2_ and, to a lesser degree, in PI(3,4)P_2_ into PI(4)P ^1,10,11^. PI3KI phosphorylates the third position of the inositol ring of PI(4,5)P_2_ into PI(3,4,5)P_3_, thereby catalyzing the inverse reaction of PTEN. This kinase is known to control the cell energetic state and metabolism and thus playing a key role in tumorigenesis ^11^. Bryant and Mostov ^1^ reported that PI(3,4,5)P_3_ is present at the basolateral membrane, but absent in the apical part, of polarized epithelial cells. PTEN and PI3K are believed to be responsible for this difference. PTEN is present in the apical part and at the tight junctions, where it transforms PI(3,4,5)P_3_ into PI(4,5)P_2_. By contrast, PI3K is located in the basolateral part of the membrane and catalyzes the opposite reaction from PI(4,5)P_2_ to PI(3,4,5)P_3_.

##### Regulation of PTEN and PI3KI

Cell polarization is highly regulated through mechanisms involving PTEN, PI3K, PI(4,5)P_2_ and PI(3,4,5)P_3_ ^18,26,27^. We investigated to what degree high activity of PTEN (2.3e-15 mg/μm^2^) and low activity of PI3KI (6.1e-16 mg/μm^2^) are sufficient to deplete PI(3,4,5)P_3_ to about 2 molecules/μm^2^ and thereby mimic the apical membrane configuration. Conversely, we asked if low PTEN (3.9e-17 mg/μm^2^) and high PI3KI (1.5e-14 mg/μm^2^) could replicate the basolateral membrane configuration, which is rich in PI(3,4,5)P_3_ (760 molecules/μm^2^). Interestingly, model simulations readily mimicked both membrane configurations, which suggests that the model is a satisfactory approximation of the observed phenomena characterizing epithelial and basolateral membrane states (Figure 6).

**Figure 6.**
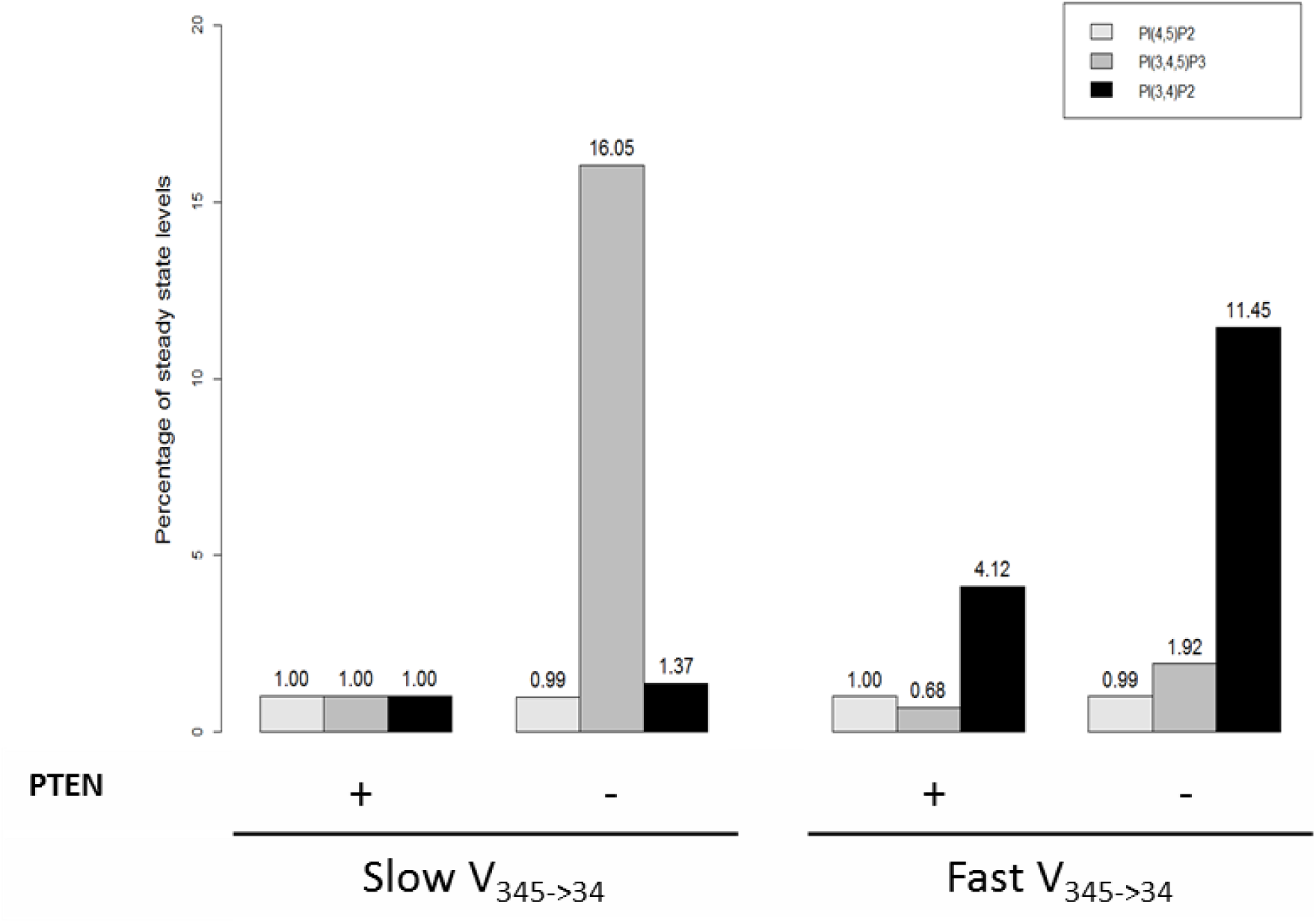
PI(3,4,5)P_3_ is sensitive to PTEN when v_345→34_ is slow. A decrease in PTEN is sufficient to increase the levels of PI(3,4,5)P_3_ and change the membrane configuration form apical (low PI(3,4,5)P_3_) to basolateral (high PI(3,4,5)P_3_). A fast v_345→34_ will decrease PI(3,4,5)P_3_ and make the membrane much less sensitive to a PTEN change. A PTEN knockdown of 98.33% does not alter the amount of PI(4,5)P_2_ in either fast or slow v_345→34_ conditions. Fast v_345→34_ increases the levels of PI(3,4)P_2_ and makes the levels of this lipid dependent on the PI(3,4,5)P_3_ pool. Slow v_345→34_ is modeled as γ_345→34_ = 1e11 and f_345→34_ = 0.9982. Fast v_345→34_ is modeled as γ_345→34_ = 6e13 and f_345→34_ = 0.9998.

##### The flux v_*345→34*_ modulates the effects of PTEN

If the flux v_345→34_ is accelerated to values close to those ones described in the literature for SH2 domain-containing phosphatidylinositol 5-phosphatase (SHIP1), the model predicts a decrease in PI(3,4,5)P_3_. The surprising consequence of this prediction is that this decrease will lock the membrane in a basal-like configuration and that a knockdown of PTEN will no longer increase PI(3,4,5)P_3_ (Figure 6).

#### 2.3.3. Control of PI(4,5)P_*2*_ levels

The proposed model is a powerful tool for exploring how the cell controls the phosphoinositide levels in its cell membrane. Due to the multiple functions of PI(4,5)P_2_, including ion channel activity regulation, cell polarization, and signaling, the control of this phosphoinositide is of particular relevance.

PI(4,5)P_2_ can be synthesized from three phosphoinositide species in addition to PI (through v_0→45_), namely PI(4)P, PI(5)P and PI(3,4,5)P_3_ (Figure 1). PI(3,4,5)P_3_ is present in low concentrations and transformed into PI(4,5)P_2_ mainly by the phosphatase PTEN. The cellular location of this enzyme is tightly regulated, as it is located in non-polarized cells in the cytosol and nucleus most of the time ^28^. PI(5)P also exists as a small pool and its role is not clearly understood. That leaves PI(4)P as the only reasonable candidate for maintaining PI(4,5)P_2_ levels, besides PI. PI(4)P is a substrate for the kinase PIP5KIγ and has a physiological concentration roughly similar to PI(4,5)P_2_ pool, *i.e.*, around 10,000 molecules/μm^2^. However, it has been observed that PI(4,5)P_2_ levels can be maintained even with low PI(4)P levels ^11,23^. Figure 3b shows the changes in PI(4)P and PI(4,5)P_2_ levels predicted by the model when different sources are perturbed.

##### Contribution of v_*0→45*_ to PI(4,5)P_*2*_ levels

The flux v_0→45_ represents the direct transformation of PI into PI(4,5)P_2_ by means of a ternary complex of proteins containing PI4K and PIP5KI ^29^. The model suggests that v_0→ v_^45^ alone can maintain 80% of the basal level of PI(4,5)P_2_, thereby making it the main source of PI(4,5)P_2_ (Figure 3b). This direct transformation of PI into PI(4,5)P_2_ should exist to ensure the stability of the PI(4,5)P_2_ pool, and reports in the literature ^29^ ^30^ seem to support this finding.

##### PI4P influx contribution to PI(4,5)P_*2*_ levels

The flux v_→4_ represents the amount of PI(4)P coming from the Golgi through vesicle trafficking or non-vesicle transfer (Figure 2), which has been reported to constitute a sizeable contribution to the maintenance of plasma membrane PI(4)P, but contributes only moderately to the maintenance of PI(4,5)P_2 31 30_. Indeed, the model simulations show that v_→4_ by itself can maintain PI(4)P at 30% of its basal level and only generates a 9% increase in the PI(4,5)P_2_ pool (Figure 3d).

##### PI5P contribution to PI(4,5)P_*2*_ levels

The flux v_5→45_ can maintain the PI(4,5)P_2_ pool at 34% of its basal level (Figure 3b). However, the influence of this flux is highly dependent on v_→3_. If _v→3_ increases 25 times, which makes this input flux similar to the one for PI(4)P, v_5→45_ can sustain PI(4,5)P_2_ levels at 71%. If v_→3_ increases 50 times, v_5→45_ can sustain 100% of PI(4,5)P_2_. This result suggests that PI(5)P may have an influential role in the maintenance of PI(4,5)P_2_ levels and function as a means of channeling material from PI(3)P toward the linear pathway of PI(4)P, PI(4,5)P_2_ and PI(3,4,5)P_3_.

Taken together, these results suggest that the cell employs at least four mechanisms to maintain adequate PI(4,5)P_2_ levels. This level of redundancy highlights the importance of PI(4,5)P_2_. Indeed, PI(4,5)P_2_ is known as a characteristic component of the cell membrane ^11^ ^23^, and it is to be expected that down-regulation of PI(4,5)P_2_ levels would interfere with the proper functioning of the proteins in the membrane. Compromising these proteins, in turn, would have a negative impact on fundamental processes, such as cellular nutrient intake, information sensing, chemical messaging and the secretion of waste.

### 2.4. Therapeutic Targets for the Modulation of ENaC Activity in CF

One of the motivations to develop this model was to explore the role of the phosphoinositide pathway in the modulation of the epithelial Na+ channel (ENaC) activity in the lung tissue of patients with CF. ENaC is a sodium and water channel whose activity is upregulated in CF. It is well established that PI(4,5)P_2_ promotes ENaC activity ^11^ ^32^. We have also previously identified the phosphoinositide pathway to be a key regulator of ENaC ^13^. Indeed, performing an siRNA screen in the CF context using a microscopy-based live-cell assay, we identified 30 enzymes in the phosphoinositide pathway as significant modulators of ENaC activity. We performed independent siRNA knockdowns of phosphoinositide enzymes and re-evaluated ENaC activity with the same live-cell assay. Assuming that if a siRNA increases PI(4,5)P_2_ it will enhance ENaC activity, we compared ENaC activity results (Supplementary Table ST10) with model predictions of an siRNA effect on PI(4,5)P_2_.

Our model predictions are consistent with four out of five siRNA assays targeting phosphoinositide kinases. As these assays were not used to calibrate model parameters, this agreement of model predictions with experimental observations supports the validity of our model. Furthermore, model simulations allow us to check if the tested siRNA perturbations may have undesirable side effects on the steady-state profile of the pathway, which were not observable in the original experiments. The results for specific pathway perturbations are discussed in the next sections.

#### 2.4.1. PIP5KI

The most direct and effective way to decrease PI(4,5)P_2_ levels is by decreasing PIP5KI (E_4→45_ and E_0→45_) or enhancing the 5-phosphatases of the SIOSS enzyme group that hydrolyze the fifth position of PI(4,5)P_2_ (E_45→4_). It is documented in the literature that decreasing PIP5KI will significantly affect PI(4,5)P_2_ levels _24_. The model predicts that a knock-out of PIP5KI will trigger a decrease in PI(4,5)P_2_ to 13% of its basal steady state (Figure 7). The performed siRNAs validation tests corroborate the model prediction (Supplementary Table ST10).

**Figure 7.**
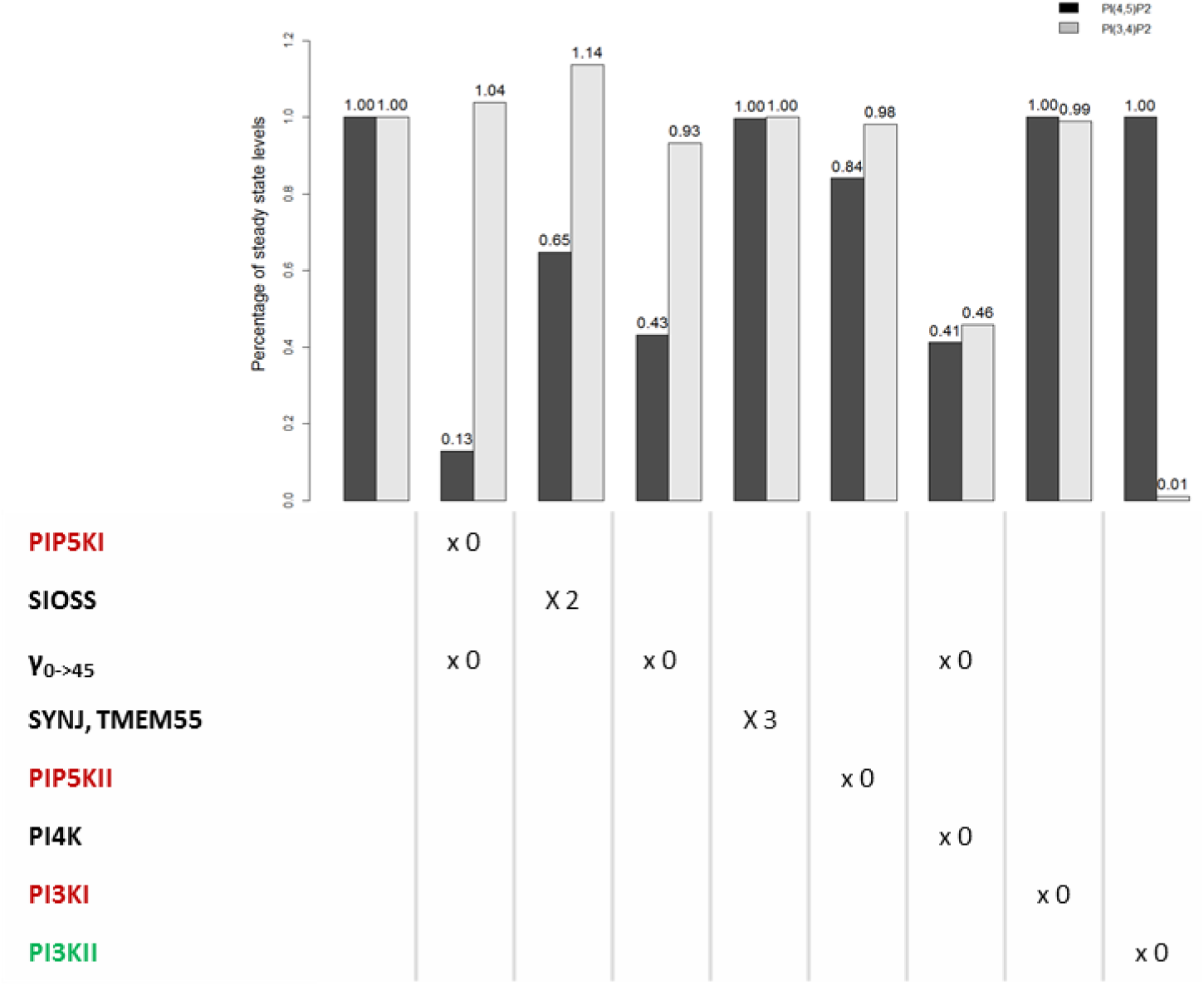
Predicted changes in the PI(4,5)P_2_ and PI(3,4)P_2_ levels as a consequence of siRNA knockdown assays. Each kinase is inactivated one at the time and phosphatases are upregulated. Because the protein complex catalyzing v_0→45_ is composed of PIP5KI and PI4K, when one of these enzymes is knocked out, the complex should be also knocked out. There is also the possibility of knocking out only the complex. Enzymes are colored according to the classification in the siRNA screens: enzymes activating ENaC are marked red, those inhibiting ENaC are marked green and those exhibiting both effects are marked black.

An alternative to trigger the decrease in PI(4,5)P_2_ levels is to increase the activity of 5-phosphatases in the SIOSS enzyme group. Doubling the activity of this phosphatase group in the model results in a 35% decrease in PI(4,5)P_2_ (Figure 7). Both our previous dataset ^13^ and results from the new siRNAs validation tests included in the present study (Supplementary Table ST10) are not as conclusive about the SIOSS phosphatases, as for most of the phosphatases tested, which could be a consequence of the unspecific activity that characterizes phosphatases. For example, synaptojanins catalyze several reactions in the pathway and perturbing them would probably cause unexpected side effects.

#### 2.4.2. PI4K

A PI4K knockout affects the fluxes v_0→4_ and v_0→45_. It decreases PI(4,5)P_2_ in 59% (Figure 7). Accordingly, the model suggests that PI4K should be classified as an ENaC activating gene, which is in line with our previous observations ^13^. An undesirable side effect within these model predictions is the change of the PI(3,4)P_2_ concentration, a lipid involved in clathrin-coated vesicle formation and activation of AKT. According to the model, this perturbation would cause a 54% decrease in the level of PI(3,4)P_2_.

#### 2.4.3. PI4K, PIP5KI and DVL protein complex

A knockout of the protein complex formed by PI4K, PIP5KI and DVL (PI4K+PIP5KI+DVL, E_0→45_), which transforms PI directly into PI(4,5)P_2_, causes a 57% decrease in this lipid (Figure 7). This perturbation also causes a 7% decrease in the levels of PI(3,4)P_2_. The PI4K+PIP5KI+DVL protein complex is formed upon wingless-Type MMTV Integration Site Family, Member 3A (Wnt3a) stimulation. The possibility of targeting the segment polarity protein Dishevelled homolog DVL (DVL) to suppress the formation of the protein complex is interesting because it would avoid interfering with other reactions in the pathway.

#### 2.4.4. PIP5KII and SYNJ/TMEM55

An increase of SYNJ/TMEM55 (E_45→5_) phosphatases and a decrease of the kinase PIP5KII (E_5→45_) could decrease PI(4,5)P_2_ (Figure 7). The model predicts that SYNJ/TMEM55 has a negligible effect, which is consistent with the literature ^11^ and our previous data ^13^ concerning synaptojanins. However, no phosphatases belonging to the TMEM55 group were screened in the Almaça *et al.* study. Knocking out PIP5KII reduces the pool of PI(4,5)P_2_ by 16%. PIP5KII was classified as an ENaC enhancer in both our previous screens ^13^ and in the present siRNA validation tests (Supplementary Table ST10), in agreement with the model prediction. Altering PIP5KII function only causes a 2% decrease in PI(3,4)P2, however perturbing PIP5KII activities could have unforeseen consequences since the role of PI(5)P is not clearly understood and the flux catalyzed by PIP5KII, v_5→45_, is the main efflux for the PI(5)P pool.

#### 2.4.5. PI3KI and PI3KII

The model predicts that PI(4,5)P_2_ levels are insensitive to knockouts of PI3KI and PI3KII. We previously classified PI3KI as an ENaC activating gene ^13^, and this is corroborated here by the siRNA validation tests. The PI3KI knockdown increases the level of PI(4,5)P_2_ if the model parameters are configured to reproduce a basolateral-like membrane composition (enriched in PI(3,4,5)P_3_). At the same time, this simulated PI3KI knockdown decreases PI(3,4,5)P_3_ levels, which is also known to control ENaC ^11^. One should note that in polarized cells ENaC localizes to the apical part of the membrane which contains neither PI(3,4,5)P_3_ nor PI3KI. Therefore, the effect of PI3KI on ENaC may only be observable in non-polarized cells.

The model predicts a negligible influence of PI3KII on PI(4,5)P_2_ and PI(3,4,5)P_3_ but causes an almost complete depletion of PI(3,4)P_2_. We previously ^13^ classified PI3KII as an ENaC inhibiting gene. If this is so, the model suggests that this inhibition could be caused by the depletion of PI(3,4)P_2_ or components not belonging to the phosphoinositide pathway.

## 3. DISCUSSION

In this work, we developed a new mathematical model that captures the complex metabolic network of phosphoinositides. The proposed model successfully replicates the phosphoinositide metabolite levels in mammalian cells and reflects numerous observed phenomena. The model is also able to reproduce the differentiation of the cell membrane into apical and basolateral types.

Using model simulations we were able to dissect the control of the levels of PI(4,5)P_2_, for which the low abundant phosphoinositide PI(5)P seems to have a significant role as an alternative source. This finding was not detectable in previous models of the pathway, due to their simplifying assumptions.

Moreover, the model was helpful in explaining observed effects in a siRNA screen of ENaC modulators in CF. Specifically, the model suggested targeting the enzyme that catalyzes v_4→45_, PIP5KI, as the most effective way to decrease the levels of PI(4,5)P_2_. Targeting PI4K would also reduce PI(4,5)P_2_ levels significantly, but model simulations point to a possible undesired side effect, namely, the simultaneous reduction of PI(3,4)P_2_ levels. Targeting the PI4K+PIP5KI+DVL protein complex does not significantly alter other lipids (Supplementary Figure S2) while yielding a large PI(4,5)P_2_ reduction. Because this reduction is not as extensive as the one induced by PIP5KI targeting, it may moderate ENaC activity without drastic negative effects in the activity of other proteins regulated by PI(4,5)P_2_.

The model also suggests that, in order to replicate phenomena retrieved from the literature, v_0→45_ should be the main flux producing PI(4,5)P_2_. In particular, this flux may explain the maintenance of PI(4,5)P2 levels when the levels of PI(4)P are low. This result suggests the importance of a close functional relationship between PI4K and PIP5KI. This relationship does not imply that the two kinases must be in physical proximity through this particular protein complex ^29^. They may also work in close proximity within lipid raft-like structures, for example.

The coupling of PI4K and PIP5KI activities may define two configurations of the system. One, where the two kinases are working together closely, in which case they are more sensitive to alterations in PI and in the levels of PI4K. The other configuration is more robust in terms of PI(4,5)P_2_ levels, where the bulk of this phosphoinositide is created through PI(4)P.

Of course, the model could be improved in the future when new experimental data regarding phosphatases and higher parameter precision are available, but this information is much scarcer than that of kinases. For instance, it is unclear how exactly phosphatases act on the system. Their versatility may suggest the existence of competitive inhibition among their substrates, but this competition could cause substrate coupling, when all substrates of a phosphatase are influenced by the alteration of a single substrate, especially if the phosphatase is saturated ^33^.

Along the same lines, some kinases catalyze multiple reactions. It would thus make sense to consider substrate competition at a more general level. In preliminary studies, we already considered substrate competition, but did not detect significant differences in model behavior.

Although the model is quite robust, it has few shortcomings. For instance, we estimated values for the parameters γ_5→45_, f_5→45_, γ_3→35_ and γ_35→5_, which differed somewhat from literature reports, in order to replicate the levels of PI(5)P and PI(3,5)P_2_. Also, not all phenomena were fully replicated by the model: PI(4,5)P_2_ did not decrease proportionally to PI, and when MTMR was reduced to 65% (simulating the knockout of MTMR2), PI(5)P did not drop to 20%, only to 98.97%. These discrepancies could be due to information gaps in the information about the system. In particular, most of the quantitative data address the total cell and are not membrane specific. Also, *in vitro* experimental results used to parameterize the model may not truly replicate the system behavior *in vivo*.

The current model does not incorporate some regulatory mechanisms which nevertheless may be implemented in future versions. For instance, Bulley *et al.* ^34^ report the activation of PTEN and PI3KI by their own products, as well as activation of myotubularins and PTEN, and inhibition of SHIP by PI(5)P.

Finally, because the phosphoinositide pathway acts differently in different organelle membranes ^2^, it could be interesting to model not only a cell membrane patch but also the membranes of the Golgi, nucleus and the endoplasmic reticulum with a multiple compartment model featuring lipid transport between them (Figure 2).

In spite of these simplifications, the proposed model is the first to successfully replicate phosphoinositide metabolism in the mammalian cell membranes. In contrast to earlier models, the model accounts for all known phosphoinositide species and permits unprecedented explorations of the roles of those phosphoinositide’s that are physiologically present in small amounts.

The model was used to identify the best approaches to control PI(4,5)P2 levels with the goal of establishing new therapeutic targets in the context of CF. The model suggests that the most effective way to accomplish this goal is to decrease the activity of the enzyme PIP5KI (v_4→45_). Additionally, v_0→45_ was also found to be very important in the maintenance of PI(4,5)P2 levels. Targeting proteins that are part of the protein complex of PI4K, PIP5KI and DVL or contribute to its control should offer an effective way to control PI(4,5)P2 levels.

In this work, we tried to arrange the current knowledge on the phosphoinositide pathway into a coherent structure. This is an important tool into the understanding of a complex layer of cell regulation that is usually overlooked and can impact fields of study with great potential to improve the human well-being like CF and cancer.

## 4. METHODS

### 4.1. Model Equations

A dynamical model of phosphoinositide metabolism was designed within the framework of Biochemical Systems Theory (BST)^21,35^–^39^, using ordinary differential equations (ODEs) in the format of a generalized mass action (GMA) system. In this approach, each ODE describes the dynamics of a dependent variable *X*_*i*_, which is formulated as a sum of all fluxes that are directly related to this variable; furthermore, each flux v_i→j_ is formulated as a power law function, as indicated in equation (1).

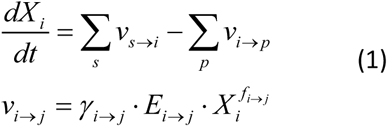

The dependent variables (*X*_*i*_) represent the actual numbers of phosphoinositide molecules (*X*_*3*_: PI(3)P; *X*_*4*_: PI(4)P; *X*_*5*_: PI(5)P; *X*_*34*_: PI(3,4)P2; *X*_*35*_: PI(3,5)P2; *X*_*45*_: PI(4,5)P2; and *X*_*345*_: PI(3,4,5)P3), and of PI (X_0_: PI) in a membrane patch of size 1μm^2^. If more than one substrate contributes to the reaction, or if the reaction is modulated by other variables, the flux term in (1) contains these contributors as additional X’s with their own powers.

The model accounts for fluxes transporting PI (v_→0_), PI(4)P (v_→4_) and PI(3)P (v_→3_) into the membrane from the ER, Golgi and endosome, respectively. Additionally, all included species were allowed to be transported out of the membrane via fluxes v_i→_. We assume that these effluxes follow first-order kinetics (f_i→_= 1) and share one common rate constant. The input flux values were restricted in order to allow 4.5% of the membrane phosphoinositides to recycle per minute (see *Supplementary Information*).

### 4.2. Parameter Estimation

Rate constants (γ_i→j_) and kinetic orders (f_i→j_) were derived from enzyme kinetic parameters obtained in BRENDA or in the literature, as detailed in the *Supplementary Information*. Enzyme activities (E i→j) and transport fluxes where manually set to approximate reported phosphoinositide steady-state values. This manually adjusted parameter set was used as an initial input for a genetic algorithm (detailed in *Supplemental Information*). This algorithm found a parameter set that minimized the deviations between model predictions and experimental observations and computing an adjustment score. The parameterized model was characterized through sensitivity and identifiability analysis, and the parameter space was explored with a Monte-Carlo approach (see *Supplemental Information*).

### 4.3. Model Implementation

The model was implemented in the programming language R v3.1.0 ^40^ together with the package deSolve ^41^. We used the ODE integration function with the LSODA method.

### 4.4. siRNA knockdown validity test

To confirm model predictions, selected phosphoinositide pathway hits identified in a large scale siRNA screen^13^ were validated with an independent round of siRNA knockdown assays. Human alveolar type II epithelial A549 cells (ATCC, Cat no. CCL-185) were transfected with 2 or 3 different siRNAs targeting phosphoinositide pathway enzymes. After transfection the FMP/Amiloride live-cell assay^13^ was applied to measure ENaC activity. Detailed methods and analysis are described in the *Supplementary Information*.

### 4.5. Data Availability

All data generated or analyzed during this study are included in this published article. Please see Supplementary Table ST10 in the *Supplementary Information* file.

## 6. ACKNOWLEDGEMENTS

Work supported by UID/MULTI/04046/2013 centre grant (to BioISI) and DIFFTARGET PTDC/BIM-MEC/2131/2014 grant (to MDA), both from FCT, Portugal. DO is a recipient of a PhD fellowship from BioSys PhD programme (Ref: SFRH/BD/52486/2014) and IU of SFRH/BD/69180/2010, both from FCT, Portugal. The authors are also grateful to Luís Marques (BioISI) and to staff from EMBL, Heidelberg (Germany), from ALMF-Advanced Light Microscopy (Beate Neumann, Christian Tischer) core facility for technical assistance.

## 7. AUTHOR CONTRIBUTION STATEMENT

D.V.O., L.L.F., E.V. and F.R.P. contributed to the conception of this project and the drafting of the manuscript. D.V.O. did the literature review, retrieved results regarding phenomena that characterize the pathway, created the model and performed the analysis. I.U. and M.D.A. performed experiments. All authors reviewed the manuscript.

